# The novel spore-specific regulator SscA governs *Aspergillus* conidiogenesis

**DOI:** 10.1101/2023.05.24.542192

**Authors:** Ye-Eun Son, Jae-Hyuk Yu, Hee-Soo Park

## Abstract

A major group of fungi produces asexual spores (conidia) for propagation and infection. Despite the critical role of conidia, the underlying mechanism of spore formation, integrity, and viability is not fully elucidated. In this study, we have identified and investigated the role of the spore-specific transcription factor (TF) SscA in three representative *Aspergillus* species. Comparative transcriptomic analyses have revealed that 25 TF encoding genes showed higher mRNA levels in conidia than in hyphae in three species. Functional and transcriptomic analyses of the 25 genes have identified SscA as a key TF for conidial formation, maturation, germination, integrity, amino acid production, and secondary metabolism in *Aspergillus nidulans* conidia. Importantly, the roles of SscA are conserved in other *Aspergillus* species. Altogether, our study demonstrates that SscA is a novel spore-specific TF that governs production of intact and functional conidial formation in *Aspergillus* species.

**Importance:** Filamentous fungi produce myriads of asexual spores are main reproductive particles and act as infectious or allergenic agents. Although the serial of asexual sporogenesis is coordinated by various genetic regulators, there remain uncharacterized transcription factors in *Aspergillus*. To understand the underlying mechanism of spore formation, integrity, and viability, we have performed comparative transcriptomic analyses on three representative *Aspergillus* species and found a novel spore-specific transcription factor, SscA. SscA has a major role in conidial formation, maturation and dormancy, and germination in *Aspergillus nidulans*. Transcriptomic data indicate that SscA coordinates conidial wall integrity, amino acid production, and secondary metabolism in *A. nidulans* conidia. Furthermore, the roles of SscA are conserved in other *Aspergillus* species. Our findings that the novel SscA has broad functions in *Aspergillus* conidia will help to understand conidiogenesis of *Aspergillus* species.

## Introduction

Fungi are ubiquitous microorganisms present throughout the world and live in various nonliving organic materials or living organisms (1, 2). *Aspergillus* species are saprophytic ascomycetous filamentous fungi that are commonly found in a wide range of environmental conditions, including air, animals, and humans (3). Among them, *Aspergillus nidulans*, is a model organism that has been used in basic science research such as genetics and cell biology for more than 70 years (4, 5). While *A. nidulans* has been explored as a versatile fungal cell factory to produce organic acids and enzymes, this fungus can cause invasive aspergillosis as an opportunistic fungal pathogen in immune-deficient people. *Aspergillus flavus* is an opportunistic fungal pathogen of agricultural plants producing foods and feeds. *A. flavus* spreads in the air in the form of asexual spores (conidia) and most can produce aflatoxins, the most potent carcinogen found in nature, which cause adverse effects on human and animal health and cause economic losses worldwide(6, 7). Moreover, *A. flavus* is the second leading cause of aspergillosis infection in immunocompromised patients, often resulting in the death of the patient (8). *Aspergillus fumigatus*, the most prevalent airborne fungal pathogen, causes allergic bronchopulmonary aspergillosis (allergic reaction) and chronic pulmonary aspergillosis in humans (9, 10). The conidia of *A. fumigatus* infect the host, adapt, and proliferate by resisting host defenses. Recent studies have demonstrated the presence of *A. fumigatus* in the lung of 20%–30% of the COVID-19 patients with, causing COVID-19-associated pulmonary aspergillosis (CAPA) with high mortality (11, 12).

Asexual development (conidiation) is the primary reproductive strategy of *Aspergillus* species. Conidia floating in the airway can germinate and proliferate under favorable conditions. The long and branching vegetative structures (hyphae) emerge as specialized foot cells extending to stalks and forming a specialized asexual sporulation structure called conidiophore bearing thousands of conidia. On the tip of the phialide, radiating conidia transform from an immature state to a mature state by modulating the cellular and physiological properties of conidia. In conidial maturation, conidial trehalose biosynthesis is commonly accomplished for tolerance to external stresses, and conidial wall compositions are modified for adaptation to desiccation and various damages (13–17). Then, the maturated conidia stay in a dormant stage for a certain duration, during which the conidial viability is maintained through the delicate control of cellular activities and active blockage of conidial germination. For instance, dormant conidia maintain low-level respiratory metabolism and cease ATP-consuming cellular activities (13, 18). They also modify transcription and translation activities, and the production of cellular primary metabolites and secondary metabolites for survival and protection (19–23). When the dormant conidia encounter the appropriate environment, they germinate by altering the conidial contents and structure.

Active research about conidiation for a few decades has demonstrated that a series of growth and developmental stages are systemically and delicately regulated by a variety of transcription factors (TFs) (13). Most TFs have DNA-binding domains that can recognize specific nucleotide sequences and regulate the expression of target genes. Representatively, the temporal and spatial activation of the BrlA-AbaA-WetA central regulatory cascade causes cessation of hyphal growth and formation of conidiophores (24, 25). VelB and VosA, members of the *velvet* family, are highly expressed in conidia and play important roles in conidial maturation and dormancy in *Aspergillus* species (26–29). SclB, one of the Zn(II)_2_Cys_6_ TFs repressed by VosA, affects proper conidiation and spore viability in *A. nidulans* (30). Another Zn(II)_2_Cys_6_ TF, inhibited by VosA, ZcfA, is essential for stress tolerance and long-term viability of *A. nidulans* conidia (31). The bZIP TFs AtfA and AtfB are required for appropriate stress resistance to oxidative stresses in *A. nidulans* conidia (32, 33).

Although various genetic regulators have been investigated, there is still limited understanding on the TFs that regulate the formation, maturation, and dormancy of conidia. This study aims to discover novel TFs that are specifically expressed in asexual spores in three representative *Aspergillus* species and to investigate the functions of these novel TFs in *Aspergillus* conidia.

## Results

### Comparative transcriptomic analyses identify 25 spore-specific TFs in three *Aspergillus* species

Conidial formation and maturation require precise spatial and temporal regulation in *Aspergillus* species (34). To understand conidiogenesis in *Aspergillus*, we investigated differentially expressed genes (DEGs) between the elongated hyphae and resting conidia of three representative *Aspergillus* species using transcriptomic analyses **(Figure S1)**. We have identified 4,754, 5,471, and 4,851 DEGs in *A. nidulans* (*Ani*), *A. flavus* (*Afl*), and *A. fumigatus* (*Afu*), respectively. Analysis employing the transcript expression profiles of orthologous groups of genes (orthogroups) (35) has revealed that 1,030 genes were commonly up- and down-regulated in the conidia of three *Aspergillus* species. We further performed gene ontology (GO) enrichment analyses and found that the up-regulated genes in conidia were significantly enriched in heterocycle biosynthesis, transcription, small molecule catabolism, and asexual spore wall assembly, and the down-regulated genes were associated with the processes of anatomical structure development, reproduction, cell differentiation, cell cycle, and fungal-type cell wall organization **(Figure 1A and Table S1)**. These findings are consistent with previous studies showing that hyphae repeatedly reproduced for vegetative filamentous growth by modulating actin filaments and cytoskeletons and differentiated into asexual structures by mediating cell integrity and cell cycle associated proteins (36). Moreover, conidia are transcriptionally active, produce various secondary metabolites, and modify the cell wall integrity for maintaining spore viability (37, 38).

**Figure 1.**
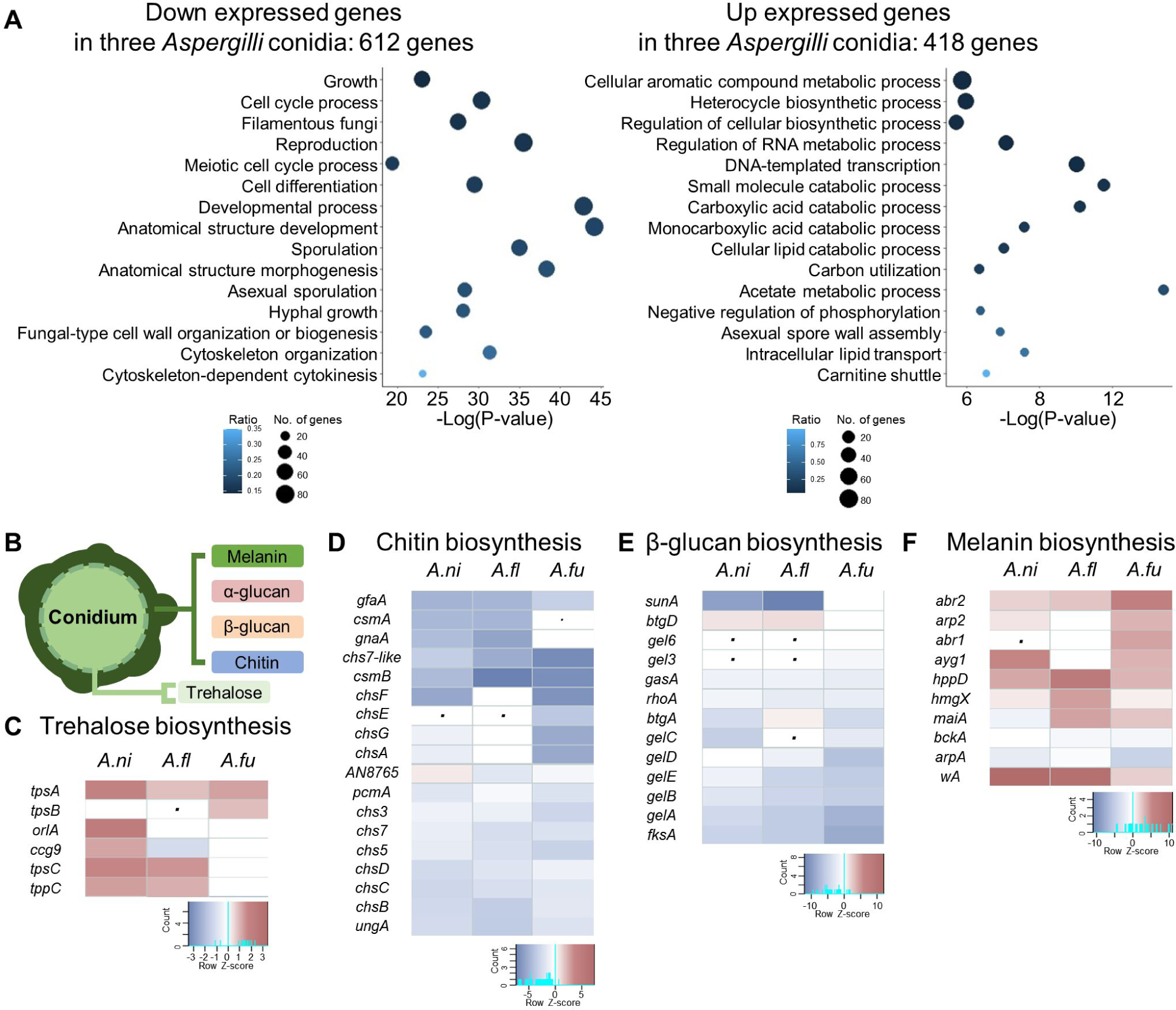
The transcriptional profiles of conidia are different from those of hyphae in three representative *Aspergillus* species. **(A)** Gene ontology (GO) enrichment analysis of commonly differentially expressed genes in the hyphae and conidia of *A. nidulans*, *A. flavus*, and *A. fumigatus*. Left: down-expressed genes in conidia, Right: up-expressed genes in conidia. **(B)** The structure of conidial wall. **(C– F)** Heat map showing the mRNA expression of genes associated with cell wall compositions. (C): Trehalose biosynthesis, (D) Chitin biosynthesis, (E) β-Glucan biosynthesis, (F) Melanin biosynthesis. The dot indicates that the corresponding homologous gene is absent.

During conidiogenesis, the cell wall components are dynamically altered. Thus, we further analyzed the expression patterns of genes contributing to the conidial structures as shown in **Figure 1B (Table 1 and Table S2)**. Although some differences existed in the three *Aspergillus* species, mRNA levels of trehalose biosynthetic genes were elevated in conidia **(Figure 1C)**. Transcript levels of chitin **(Figure 1D)** and β-glucan **(Figure 1E)** biosynthetic genes were lowered, whereas those of melanin biosynthetic genes were increased in conidia compared with those in hyphae **(Figure 1F)**. Collectively, the transcriptomic data suggest that fungal growth, development, cell wall organization, and cellular metabolism involve diverse differentially expressed genes in conidia or hyphae.

**Table 1.**
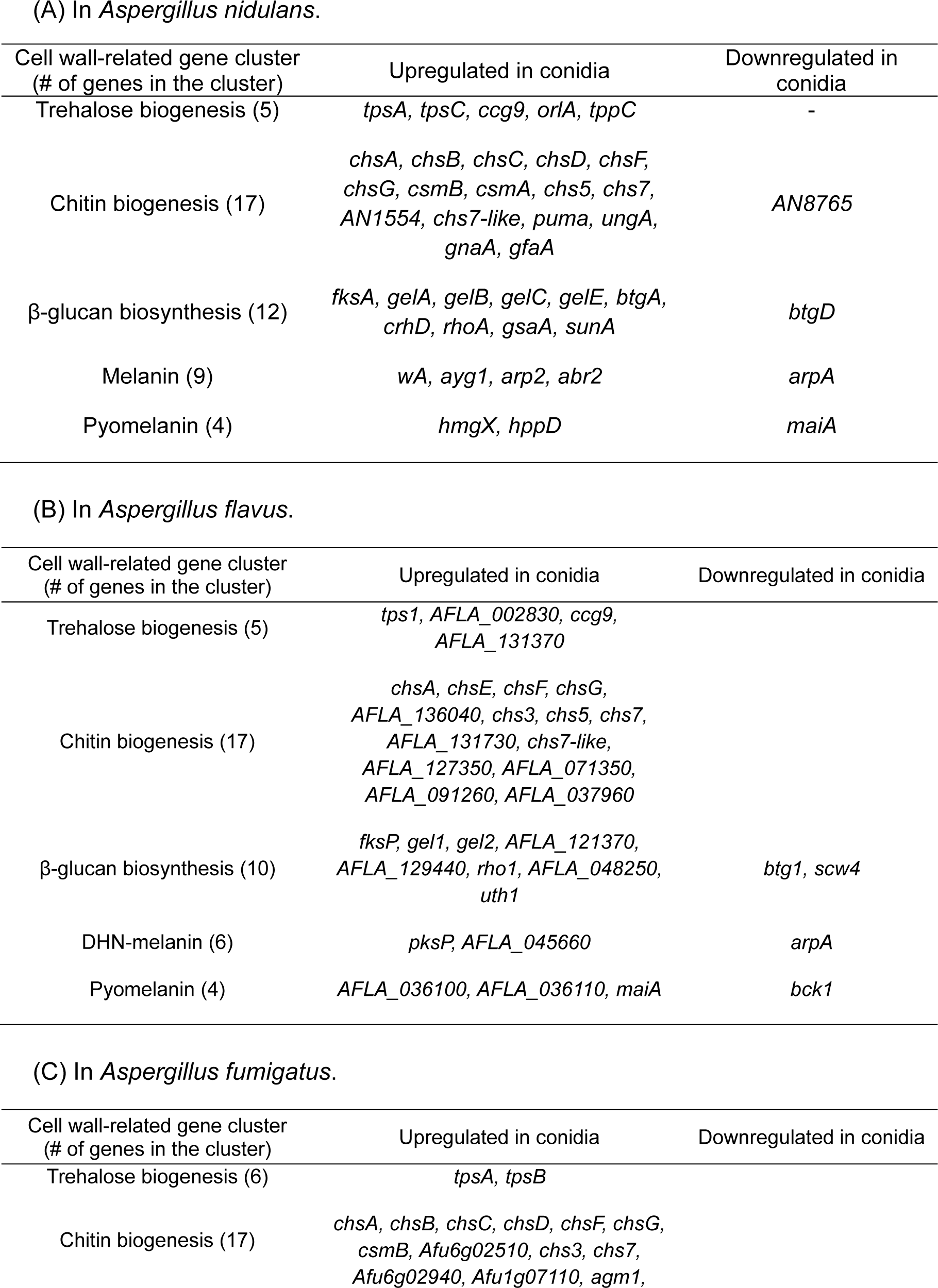
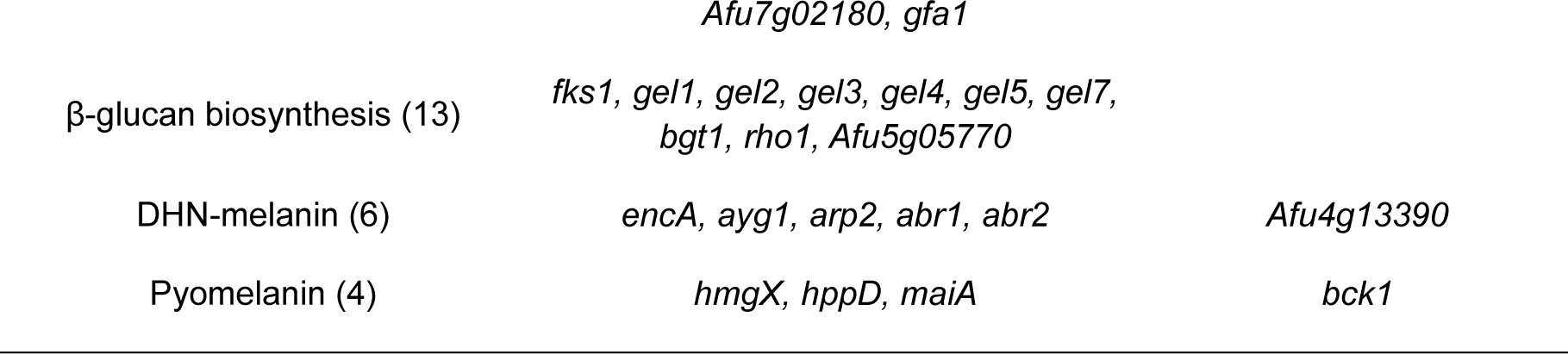
Differentially expressed genes (DEGs) associated with cell wall integrity in three *Aspergillus* species’ conidia versus hyphae.

### Functional analyses of 24 genes predicted to encode TFs in *A. nidulans*

To pin-point novel TFs regulating conidiogenesis, we analyzed the total TFs in the three *Aspergillus* species based on the protein family database (Pfam) and previous research (39, 40). We have identified 574, 576, and 492 putative TFs in *A. nidulans*, *A. flavus*, and *A. fumigatus* **(Figure S2)**, respectively, and more than half of TFs were Zn(II)_2_Cys_6_ TFs. Using transcriptomic data, we have identified significantly differently expressed TFs in the three *Aspergillus* species **(Figure 2A–2C)**. Next, we investigated commonly up-regulated TFs in the conidia of the three *Aspergillus* species. We found 25 TFs that were specifically expressed in the conidia of *Aspergillus* species **(Table 2)**, of which 19 TFs have the Zn(II)_2_Cys_6_ DNA-binding domain, termed as **<u>s</u>**pore-**<u>s</u>**pecific **<u>G</u>**al4-like zinc finger SsgA∼SsgR. Two TFs have both Zn(II)_2_Cys_6_ and C_2_H_2_ DNA-binding domains (**<u>s</u>**pore-specific **<u>C</u>**_2_H_2_ and **<u>G</u>**al4-like zinc finger ScgA and ScgB). A C_2_H_2_ TF (**<u>s</u>**pore-**<u>s</u>**pecific **<u>C</u>**_2_H_2_ zinc finger SscA) and three RING finger TFs (**<u>s</u>**pore-**<u>s</u>**pecific **<u>R</u>**ING finger SsrA∼SsrC) are also identified. One of the *velvet* family members, VosA, was confirmed in a list of up-regulated TFs in the conidia of the three *Aspergillus* species are reported in previous studies (28, 41, 42).

**Figure 2.**
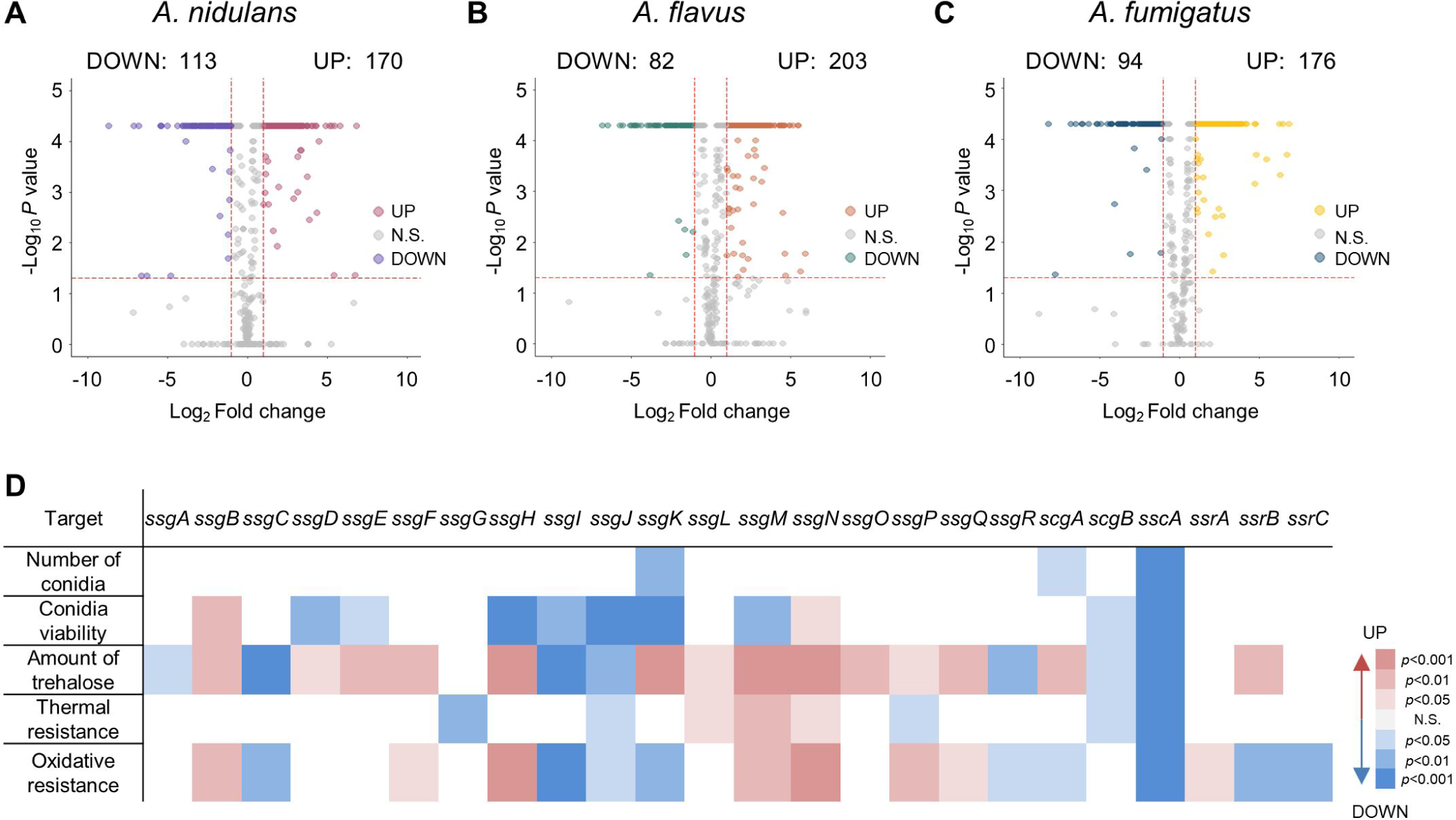
The twenty-five putative spore-specific transcription factors in three representative *Aspergillus* species. **(A–C)** Volcano plots showing up- and down-expressed TFs encoded by *A. nidulans* (A), *A. flavus* (B), and *A. fumigatus* (C). Pink, orange, and yellow points are TFs with *p* < 0.05 and Log_2_FC≥1. Ultramarine blue, green, and blue points are TFs with *p* < 0.05 and Log_2_FC≤-1. Up: up-expressed in conidia, Down: down-expressed in conidia, N.S.: not significantly expressed in conidia. **(D)** Phenome heat map showing the results of screening 24 spore-specific transcription factor deletion strains against conidiation, long-term viability, amount of trehalose, and thermal, and oxidative stress tolerance. Right color key represents reduction (blue) and enhancement (red), respectively, compared with the wild-type strain.

**Table 2.** List of transcription factors specifically up-regulated in three *Aspergillus* species’ conidia compared with hyphae.

| Domain | Gene ID |  |  | log<sub>2</sub>FC |  |  | Gene Name |
| --- | --- | --- | --- | --- | --- | --- | --- |
|  | *A. ni* | *A. fl* | *A. fu* | *A. ni* | *A. fl* | *A. fu* |  |
| **Zn(II)<sub>2</sub>Cys<sub>6</sub> (18)** | AN0742 | AFLA_070970 | Afu1g15850 | 1.28 | 1.81 | 1.07 | *ssgA* |
|  | AN1123 | AFLA_074230 | Afu1g11715 | 3.20 | 4.53 | 4.32 | *ssgB* |
|  | AN3433 | AFLA_103640 | Afu3g05760 | 3.46 | 2.46 | 2.71 | *ssgC* |
|  | AN3501 | AFLA_009580 | Afu4g14590 | 2.04 | 4.62 | 1.49 | *ssgD* |
|  | AN3502 | AFLA_107030 | Afu1g06540 | 2.48 | 1.99 | 1.43 | *ssgE* |
|  | AN3768 | AFLA_096100 | Afu3g03920 | 2.16 | 2.47 | 2.26 | *ssgF* |
|  | AN4821 | AFLA_101970 | Afu8g01150 | 1.80 | 1.30 | 3.08 | *ssgG* |
|  | AN5259 | AFLA_025470 | Afu3g02920 | 4.12 | 3.99 | 2.90 | *ssgH* |
|  | AN5390 | AFLA_008530 | Afu6g13770 | 2.35 | 2.64 | 2.02 | *ssgI* |
|  | AN6747 | AFLA_096330 | Afu7g00652 | 5.17 | 3.64 | 3.99 | *ssgJ* |
|  | AN7508 | AFLA_129420 | Afu2g05310 | 2.44 | 3.82 | 1.26 | *ssgK* |
|  | AN8079 | AFLA_024740 | Afu5g01700 | 2.31 | 4.32 | 1.18 | *ssgL* |
|  | AN9117 | AFLA_071050 | Afu7g01890 | 2.33 | 1.51 | 1.27 | *ssgM* |
|  | AN10548 | AFLA_113510 | Afu4g06420 | 1.89 | 3.04 | 1.28 | *ssgN* |
|  | AN11003 | AFLA_062330 | Afu8g07360 | 1.82 | 3.36 | 3.67 | *ssgO* |
|  | AN11112 | AFLA_131790 | Afu6g03010 | 2.40 | 4.96 | 1.01 | *ssgP* |
|  | AN11185 | AFLA_070970 | Afu7g01810 | 1.60 | 1.81 | 2.98 | *ssgQ* |
|  | AN12148 | AFLA_064980 | Afu5g02690 | 2.07 | 4.66 | 2.70 | *ssgR* |
| **C<sub>2</sub>H<sub>2</sub> & Zn(II)<sub>2</sub>Cys<sub>6</sub> (2)** | AN0096 | AFLA_093070 | Afu5g12060 | 2.36 | 4.03 | 1.06 | *scgA* |
|  | AN4013 | AFLA_028760 | Afu1g04110 | 2.5 | 3.45 | 1.70 | *scgB* |
| **C<sub>2</sub>H<sub>2</sub> (1)** | AN5003 | AFLA_084970 | Afu3g09820 | 1.13 | 1.10 | 1.67 | *sscA* |
| **Zf_RING (3)** | AN0441 | AFLA_029330 | Afu1g04600 | 5.46 | 3.81 | 1.92 | *ssrA* |
|  | AN1075 | AFLA_067440 | Afu1g12080 | 1.59 | 1.50 | 3.17 | *ssrB* |
|  | AN7713 | AFLA_062060 | Afu5g08230 | 2.12 | 1.27 | 2.19 | *ssrC* |
| **Velvet (1)** | AN1959 | AFLA_026900 | Afu4g10860 | 3.73 | 2.12 | 3.12 | *vosA* |

To explore the functions of the novel 24 TFs in *A. nidulans*, excluding the previously studied VosA, we first confirmed the mRNA levels of 24 genes. As shown in **Figure S3**, all the 24 genes were up-regulated in the last stage of asexual development and the conidia of *A. nidulans*. We then generated individual null mutant for each of the novel 24 genes in *A. nidulans* (43) and examined the phenotypes of each null (Δ) mutant (**Figure S4**). Distinct from other 23 mutants, the *sscA* null mutant exhibited unusual colored colonies in *A. nidulans*. To further understand the functions of each spore-specific TFs in conidia, we examined all null mutant for conidiation, conidial viability, conidial trehalose content, and conidial stress tolerance under thermal and oxidative stresses. Although other strains exhibited slightly different phenotypes compared with wild-type (WT), the deletion of *sscA* substantially affected all the conidial phenotypes **(Figure 2D)**. These findings led us to conduct in-depth studies on the biological roles of SscA in *A. nidulans* conidia.

### Key role of SscA in conidial formation

SscA was highly conserved in most Ascomycota **(Figure 3A)**, and the domain analysis of SscA in *Aspergillus* species showed that the C_2_H_2_ zinc finger domain was conserved in the N-terminal end **(Figure 3B)**. To confirm the effects of *sscA* deletion on asexual development, WT, Δ*sscA*, and *sscA*-complemented (Cʹ *sscA*) strains were point-inoculated and incubated. As shown in **Figure 3C**, the Δ*sscA* mutant exhibited apricot-colored fungal growth and small and incomplete conidiophores compared with those of WT and Cʹ *sscA* strains. Quantitative analyses revealed that the Δ*sscA* mutant had considerably reduced conidial production in *A. nidulans* **(Figure 3D)**. These findings suggest that SscA plays a vital role in conidial formation in *A. nidulans*.

**Figure 3.**
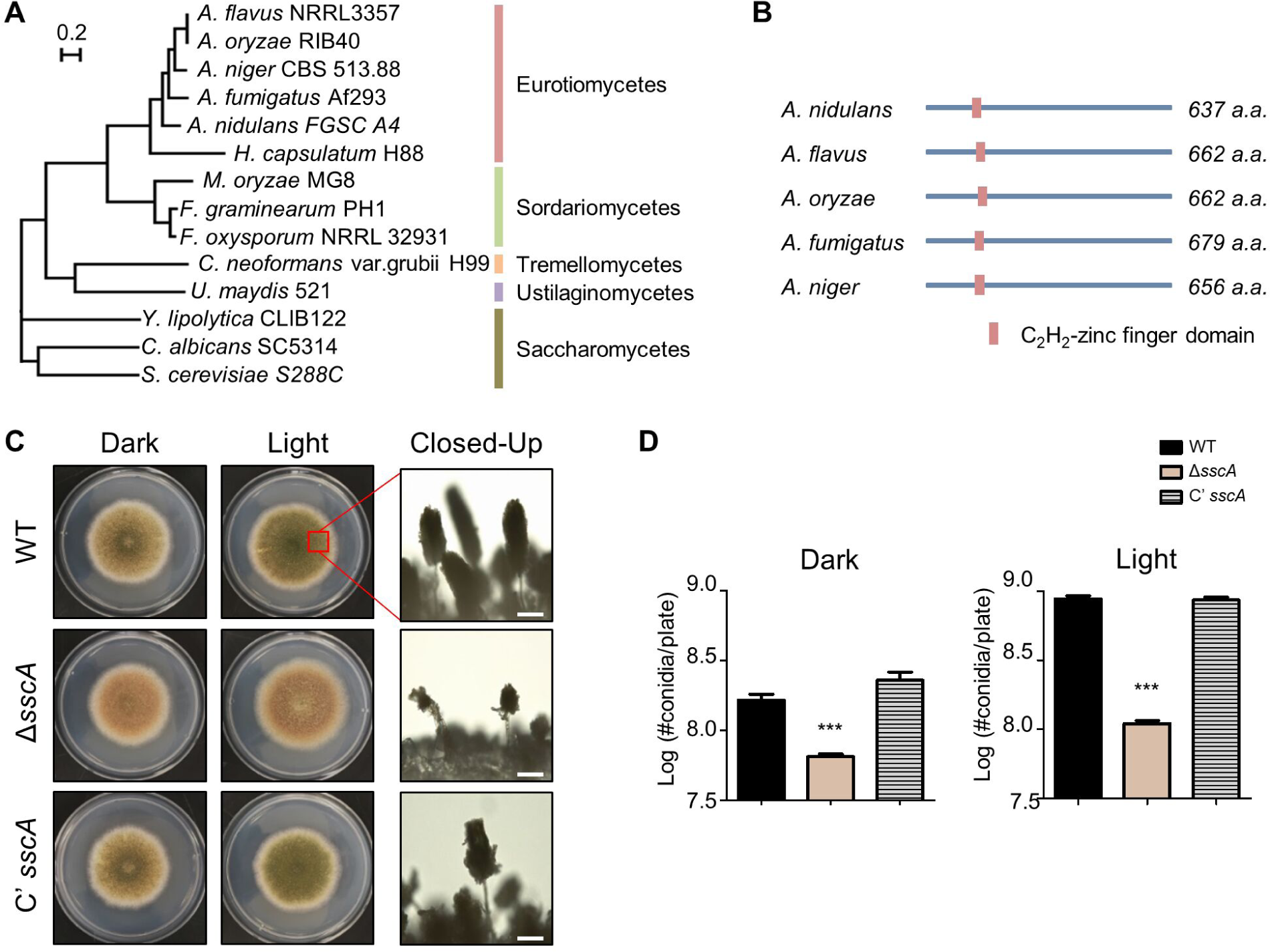
The conserved C_2_H_2_ protein, SscA, is important for asexual development and cxonidial formation. **(A)** Phylogenetic tree of SscA homolog proteins in fungi. Protein sequences were aligned with Clustal W and Mega X software, using the maximum likelihood method based on the Jones-Taylor-Thornton model with 1000 bootstrap replicates. **(B)** Domain architecture of the SscA homologs in *Aspergillus* species. **(C)** Growth phenotypes of wild-type (WT), Δ*sscA*, and complementary (Cʹ *sscA*) strains grown on solid minimal media (MM) at 37°C for 5 days under light and dark conditions. Right panels show the conidiophores of each strain observed by microscopy (bar = 0.25 mm). **(D)** Quantitative examination of conidial production shown in (C). Error bars indicate standard deviations of three biological replicates (\*\*\**p* < 0.001). **(E–F)** Quantitative real-time RT-PCR analysis of *brlA* (E) or *abaA* (F) in WT, Δ*sscA*, and Cʹ *sscA* strains after inducing asexual development. The mRNA expression was normalized to that of the endogenous control *β-actin* (\*\*\**p* < 0.001, \*\**p* < 0.01, \**p* < 0.05).

### SscA is required for proper conidial dormancy and germination

As shown earlier in **Figure 2D**, Δ*sscA* caused a decrease in the amount of trehalose, which is a key spore component for stress resistance (44). To further explore the multiple roles of SscA in conidial dormancy and germination, we evaluated the long-term viability and conidial germination of the Δ*sscA* conidia. We first observed the conidia of each strain grown for 10 days by transmission electron microscopy (TEM). As shown **Figure 4A**, conidia of WT and complemented strains exhibited full cytoplasm and normal organelle structures, termed intact conidia, but the conidia of Δ*sscA* strain exhibited loss of cytoplasm and deficient organelles. The phenotype of the 10-day-old Δ*sscA* conidia closely resembled those of the Δ*vosA* mutant (27). Quantitative analysis revealed that <10% of the Δ*sscA* mutant conidia were intact **(Figure 4B)**. To determine the long-term viability of the 10-day-old Δ*sscA* conidia, we checked the ability of 10-day-old conidia to form germ tubes and found that the survival rate of the Δ*sscA* conidia was dramatically decreased compared to those of WT and complemented strains **(Figure 4C)**. Overall, these results demonstrate that SscA is required for proper conidial viability, cellular and metabolic integrity, and dormancy.

**Figure 4.**
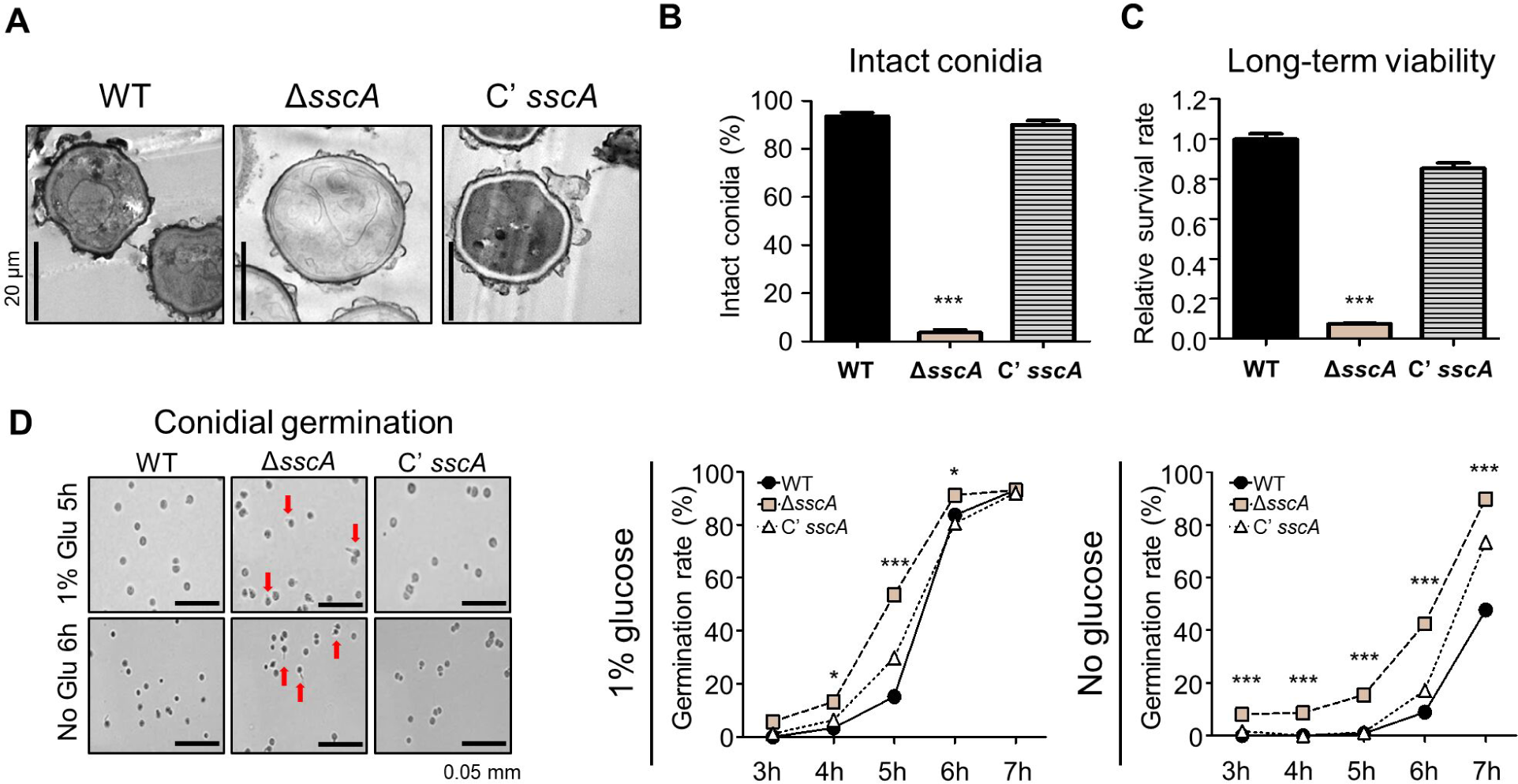
SscA is indispensable for conidial maturation, dormancy, and germination. **(A)** Transmission electron microscopic analysis of 10-day-old conidia of WT, Δ*sscA*, Cʹ *sscA* strains. The *sscA*-deleted conidia showed the loss of cytoplasm and organelles, indicating defective viability. **(B)** The percentages of intact conidia shown in (A). Intact conidia were calculated as the ratio of the number of conidia having normal cytoplasm and organelles per total conidia in 10 photographs (\*\*\**p* < 0.001). **(C)** Bar plot representing the conidial viability of WT, Δ*sscA*, and Cʹ *sscA* strains grown for 10 days (\*\*\**p* < 0.001, \*\**p* < 0.01). **(D)** Photographs of conidial germination of WT, Δ*sscA*, Cʹ *sscA* strains inoculated onto solid minimal media with or without 1% glucose after incubating for 5 or 6 h, respectively (bar = 50 μm). Right graphs represent conidial germination rate of WT, Δ*sscA*, and Cʹ *sscA* grown on MM with or without glucose source (\*\*\**p* < 0.001, \**p* < 0.05).

Next, we investigated conidial germination of the 2-day old Δ*sscA* conidia. We inoculated the fresh conidia of WT and mutant strains into agar media with or without glucose and quantified the germination rates. The germination rates of the WT, Δ*sscA*, and Cʹ *sscA* conidia were 15%, 55%, and 27%, respectively, at 5 h of incubation in the presence of 1% glucose **(Figure 4D)**. Furthermore, the germination rate of the Δ*sscA* conidia without glucose was higher than that of the WT and complemented conidia. Altogether, these results suggest that SscA is necessary for proper conidial germination in *A. nidulans*.

### The absence of *sscA* affects trehalose biosynthesis and stress response

To elucidate the molecular functions of SscA in conidia, we performed genome-wide expression analyses of the conidia of WT and Δ*sscA* strains using RNA-sequencing. Total of 4,569 genes (|Log_2_FC|≥1; *p*-value < 0.05) were differently expressed between WT and Δ*sscA* conidia **(Figure S5A)**. In the Δ*sscA* conidia, 1,874 DEGs were down-regulated and 2,695 DEGs were up-regulated **(Figure 5A)**. GO enrichment analysis of the up-regulated DEGs revealed the enrichment of genes involved in cell wall macromolecule metabolic process, glucan metabolic process, cellular amino acid catabolic process, secondary metabolite biosynthetic process, and sterigmatocystin metabolic process. GO enrichment analysis of the down-regulated DEGs showed the enrichment of genes related to response to stimulus, chemicals, and stress; gene expression; RNA metabolic process; and ncRNA metabolic process **(Figure 5A, Figure S5B, and Table S3)**. Among the DEGs affected by SscA, we correlated gene expression with phenotype. First, we found that mRNA levels of trehalose biosynthesis genes, including *tpsA*, *tpsC*, and *orlA*, were decreased in the Δ*sscA* conidia **(Figure 5B)**. Similar to the above-described screening results **(Figure 2D)**, amount of trehalose in the *sscA* null conidia was reduced compared to WT and complemented strains **(Figure 5B)**. Second, mRNA levels of genes associated with DNA repair and stress response were decreased **(Figure 5C)**, implying that the *sscA* null conidia are likely sensitive to various stresses. In fact, when the Δ*sscA* conidia were tested for the sensitivity to UV, thermal, and oxidative stresses, as depicted in **Figure 5D**, the conidia of *sscA* null mutant were more sensitive to UV, thermal, and H_2_O_2_ stresses than those of WT and complemented strains.

**Figure 5.**
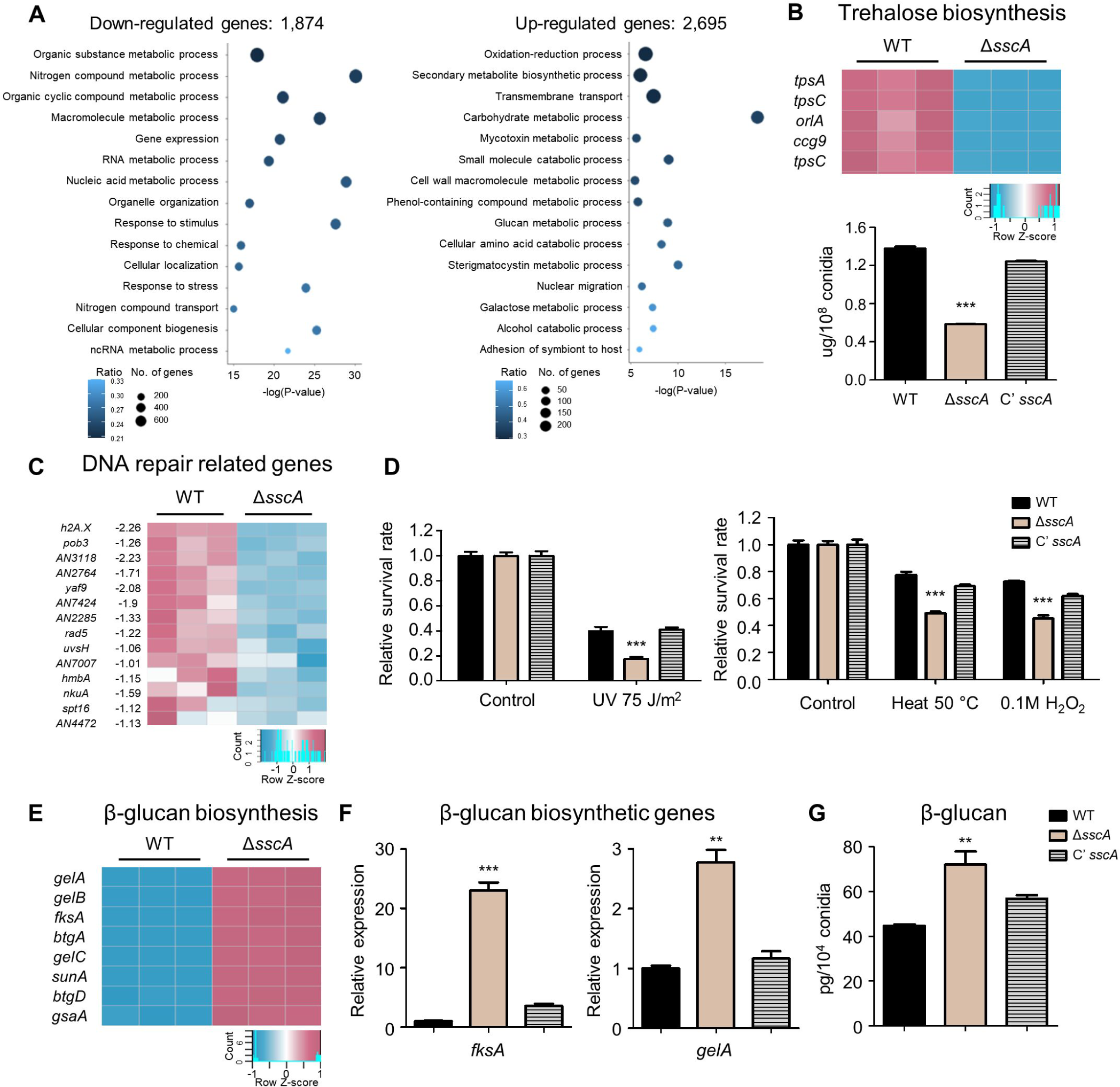
SscA is essential for stress tolerance and conidial wall integrity. **(A)** Gene ontology (GO) analysis of significantly upregulated (2,695) and downregulated (1,874) genes in the conidia of *sscA*-deleted strain compared to that in the conidia of wild-type strain (biological process). **(B)** Transcript levels of trehalose biosynthetic genes in WT and Δ*sscA* conidia. Below histograms indicate the amount (μg) of trehalose in the conidia of WT, Δ*sscA*, and Cʹ *sscA* strains grown for 2 days (\*\*\**p* < 0.001). **(C)** Transcription levels of 14 genes involved in DNA repair system, which were downregulated in the *sscA* deletion mutant. **(D)** Bar graphs showing the relative survival rate of WT, Δ*sscA*, and Cʹ *sscA* conidia stressed by ultraviolet radiation, heat, and oxidative reagents. The viable rate was calculated as the ratio of the number of survived colonies to that of the untreated control. Error bars indicate standard deviations of three biological replicates (\*\*\**p* < 0.001). **(E)** Transcription pattern of β-glucan biosynthetic genes in WT and Δ*sscA* conidia. **(F)** Bar plots showing the relative expression levels of β-glucan biosynthetic genes (*fksA* and *gelA*) in the designated strains (\*\*\**p* < 0.001, \*\**p* < 0.01). **(G)** The quantified β-glucan levels (pg) in WT, Δ*sscA*, and Cʹ *sscA* conidia (\*\**p* < 0.01).

### SscA is essential for proper conidial wall integrity in *A. nidulans*

RNA-seq data showed that mRNA levels of genes related to the cell wall macromolecule metabolic process were increased in the Δ*sscA* conidia **(Figure 5A)**. In particular, mRNA levels of genes associated with glucan and chitin biosynthesis were highly increased in the Δ*sscA* conidia **(Figure 5E)**. We confirmed this result by qRT-PCR assay and found that expression of two beta-glucan biosynthetic genes *fksA* and *gelA* was increased in the Δ*sscA* conidia **(Figure 5F)**. We then examined the amount of β-glucans in the conidia of WT, Δ*sscA*, and Cʹ *sscA* strains and asked whether the changes in gene expression direct the phenotype. As shown **Figure 5G**, the amount of β-glucan in the Δ*sscA* conidia was high compared to that in the conidia of WT and Cʹ *sscA* strains, suggesting that SscA is crucial for proper β-glucan levels in the conidia. Furthermore, the transcript levels of genes related to chitin and hydrophobin biosynthesis were up-regulated in the Δ*sscA* conidia **(Figure S5)**. Overall, these results indicate that SscA is a key regulator of conidial wall structure and integrity in *A. nidulans*.

### SscA down-regulates the production of sterigmatocystin (ST) and other secondary metabolites in *A. nidulans* conidia

In the Δ*sscA* conidia, mRNA levels of genes associated with the secondary metabolite biosynthetic process and sterigmatocystin (ST) metabolic process were increased, implying that SscA inhibits the production of various secondary metabolites in *A. nidulans* conidia. To test this hypothesis, we analyzed expression of the ST gene cluster in WT and the Δ*sscA* conidia. As shown in **Figure 6A**, the expression of most ST biosynthetic genes was increased in the Δ*sscA* conidia. The results of RNA-seq were further confirmed by qRT-PCR analysis **(Figure 6B)**. Amount of ST in the Δ*sscA* conidia was increased compared to that in the conidia of WT and complemented strains **(Figure 6C)**. Moreover, the amount of ST production in the Δ*sscA* mycelia was also increased compared to those of WT and Cʹ *sscA* strains **(Figure S6)**. To further comprehend the roles of SscA in ST biosynthesis, we generated the *sscA*-overexpressed strain (OE*sscA*) by expressing *sscA* under the *alcA* promoter. As shown in **Figure 6D**, OE*sscA* strain produces much less amount of ST than the control strain under noninducing as well as inducing conditions. These data suggest that SscA act as a repressor of ST biosynthesis. We also checked expression patterns of 25 secondary metabolite gene clusters **(Table S4)**. In particular, the transcript levels of asperthecin, emericellamide, and terrequinone biosynthetic genes were all elevated in the Δ*sscA* conidia compared to WT conidia **(Figure S7)**. Altogether, these results suggest that SscA plays as a master-controller for the production of various secondary metabolites in *A. nidulans* conidia.

**Figure 6.**
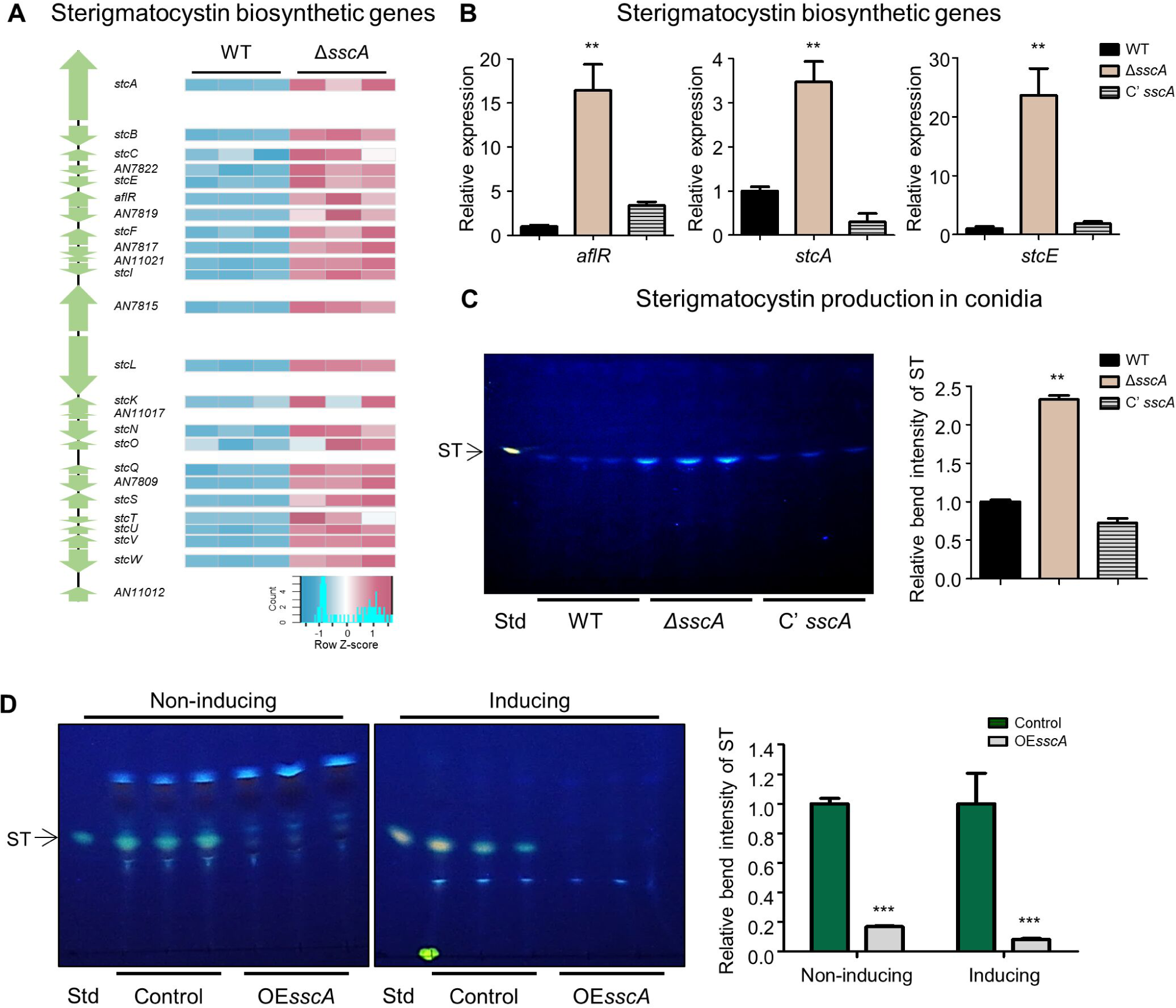
SscA contributes to sterigmatocystin biosynthesis process in conidia. **(A)** A diagram of sterigmatocystin (ST) biosynthesis gene cluster. Right heatmap shows the relative transcript abundance of the ST gene cluster. **(B)** Bar plots indicate the mRNA expression levels of ST biosynthetic genes (*aflR*, *stcA*, and *stcE*) in the designated strains (\*\**p* < 0.01). **(C)** Thin-layer chromatography (TLC) image of ST extracted from WT, Δ*sscA*, and Cʹ *sscA* conidia. Right bar plot shows the relative production of ST analyzed in (B) (*\*\*p* < 0.01). **(D)** TLC analysis of sterigmatocystin (ST) produced by control and OE*sscA* under liquid non-inducing and inducing media. The right bar graph represented the relative bend intensity of ST (\*\*\**p*<0.001).

### SscA contributes to the regulation of amino acid catabolism in *A. nidulans* **conidia**

Transcriptomic data revealed that mRNA levels of amino acid catabolism genes were altered in the Δ*sscA* conidia. The amino acid biogenesis pathway is controlled by various molecular regulators, and SscA affected mRNA levels of those genes **(Figure 7A)**. To corroborate this, we evaluated the levels of amino acids in WT conidia and the Δ*sscA* conidia. The Δ*sscA* conidia exhibited considerably diminished levels of free amino acids, excluding leucine, compared to those of WT conidia. Especially, amounts of three basic amino acids were markedly decreased in the Δ*sscA* conidia **(Figure 7B)**. Collectively, these results indicate that SscA governs appropriate amino acid biosynthesis in *A. nidulans* conidia.

**Figure 7.**
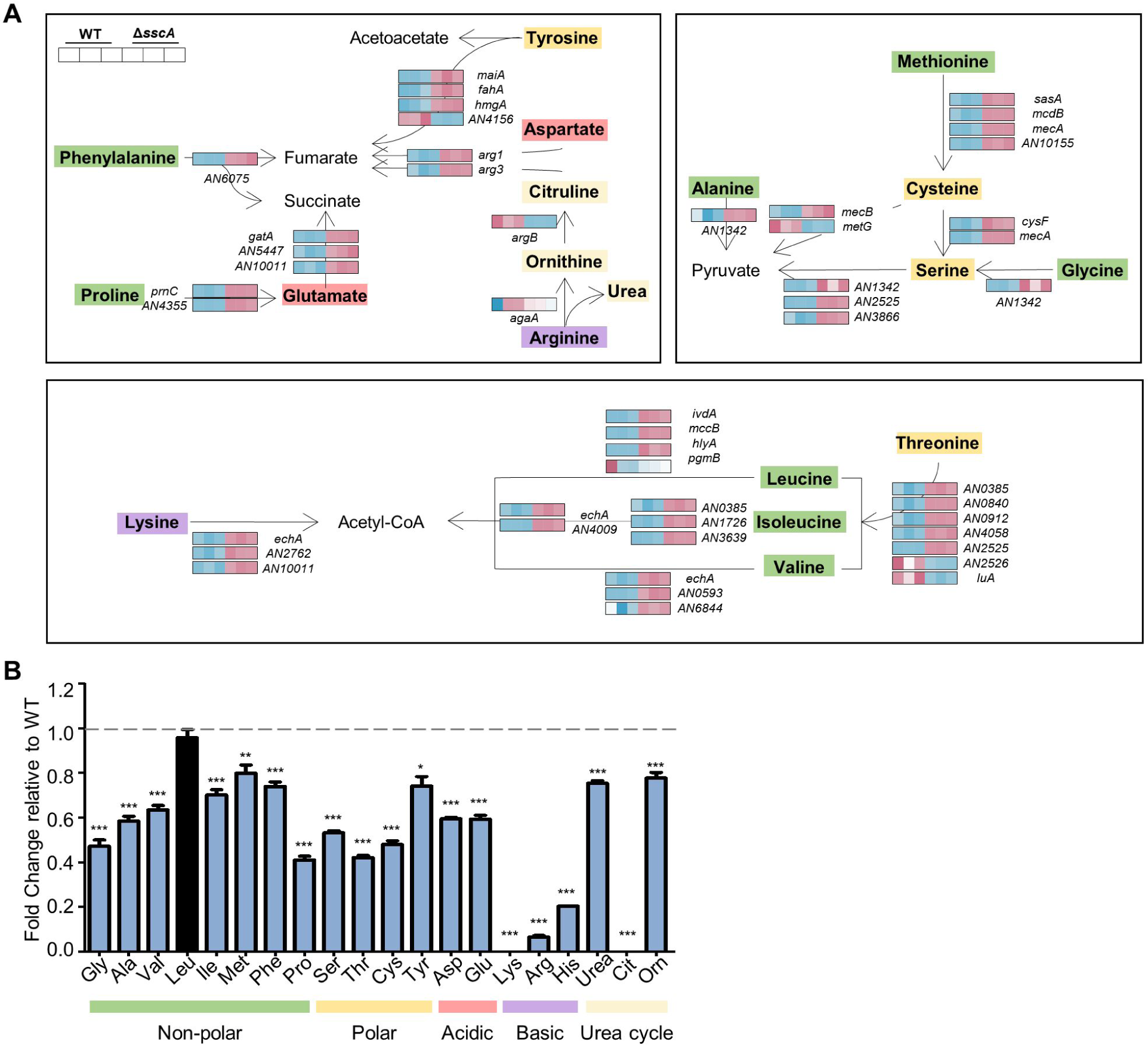
SscA contributes to the regulation of primary metabolites in conidia. **(A)** Heatmap plot representing significantly differentially expressed genes involved in amino acid catabolism in *A. nidulans*. **(B)** Free amino acid profiling of wild-type and Δ*sscA* conidia using the amino acid analyzer. The relative levels were calculated as the amount of amino acids in Δ*sscA* conidia to that of amino acids in WT conidia (\*\*\**p* < 0.001, \*\**p* < 0.01, \**p* < 0.05).

### The role of SscA is conserved in the genus *Aspergillus*

As mentioned earlier, mRNA level of *sscA* were high in the conidia of three representative *Aspergillus* spp.. To assess whether the role of SscA is conserved in the three *Aspergillus* species, we generated complemented strains that expressed *AflsscA* or *AfusscA* in the background of *AnisscA* deletion. As shown in **Figure 8**, the changes in colony color and the number of conidia affected by the absence of *AnisscA* were fully restored back to WT phenotype by heterologously expressing *AflsscA* or *AfusscA*. The decrease in the amount of trehalose and stress sensitivity caused by *AnisscA* deletion was also restored to WT levels in *AflsscA*- or *AfusscA*-expressed strains. These results strongly indicate that SscA plays conserved roles in conidial production, trehalose biosynthesis, and stress response in the genus *Aspergillus*.

**Figure 8.**
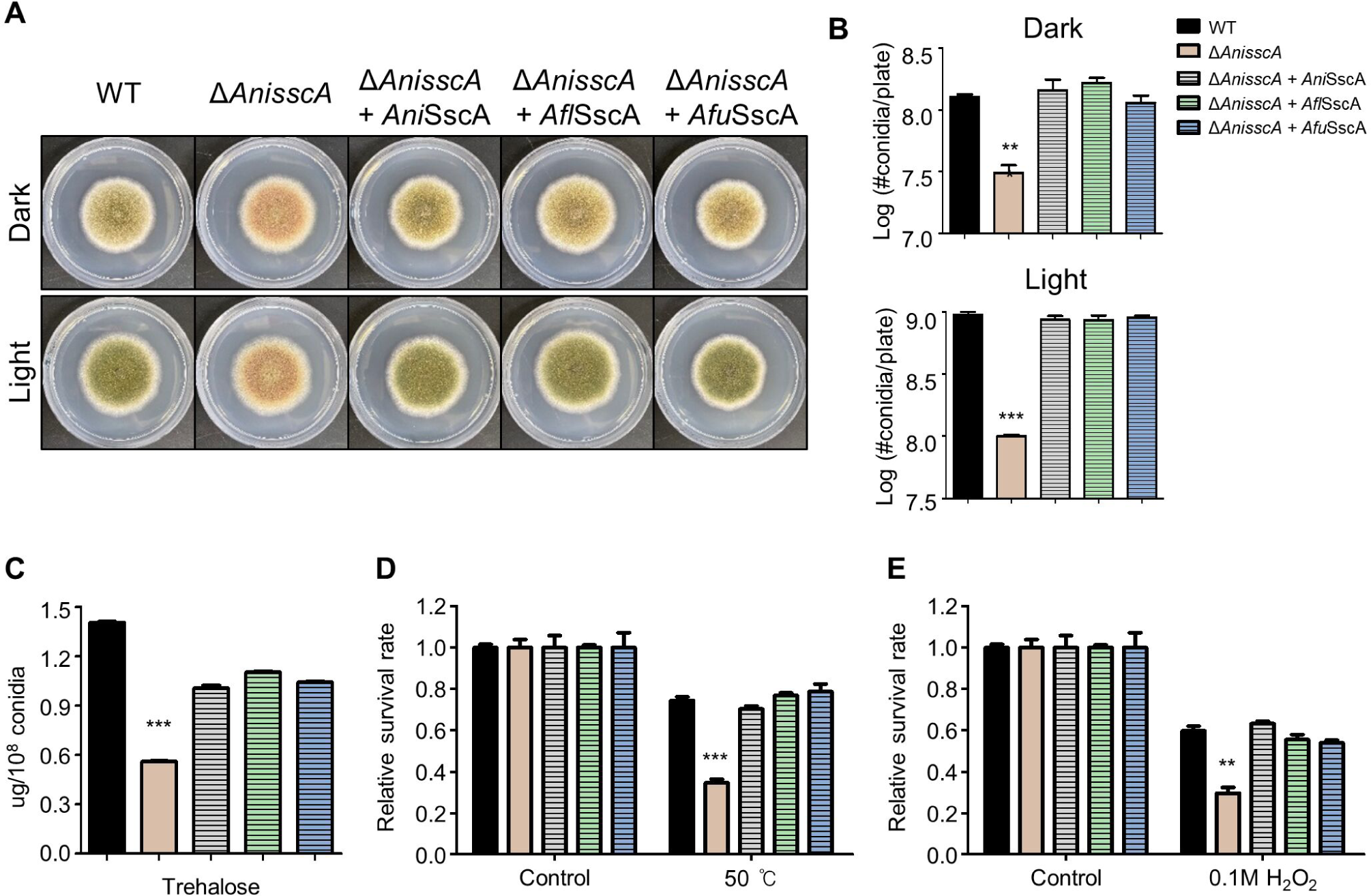
SscA of *A. flavus* and *A. fumigatus* has conserved roles in the physiology of *A. nidulans*. **(A)** Colony morphology of wild-type, ΔA*nisscA*, Cʹ *AnisscA*, Cʹ *AflsscA*, and Cʹ *AfusscA* strains grown on solid minimal media (MM) at 37°C for 5 days under light and dark conditions. **(B)** Quantitative data of conidial production shown in (A) (\*\*\**p* < 0.001, \*\**p* < 0.01). **(C)** Bar plot indicating the amount (μg) of trehalose in the conidia of the designated strains (\*\*\**p* < 0.001). **(D–E)** Histograms representing the survival rate of the conidia of the designated strains against thermal (D, 50°C) and oxidative (E, 0.1 M H_2_O_2_) stresses. The survival rates were calculated as the ratio of the number of viable colonies to that of the untreated control. Error bars indicate standard deviations of three biological replicates (\*\*\**p* < 0.001, \*\**p* < 0.01).

## Discussion

*Aspergillus* conidia, the primary propagules, are adapted for long-term viability, stress tolerance. During conidial maturation, cellular transcription and translation are activated, and the accumulated transcriptome and proteome allow resting conidia to break and resume spore germination and reproduction (13, 45–47). Although studies on *Aspergillus* conidia have been conducted for decades, there are limited functional studies on TFs that are crucial for conidial maturation and dormancy in *Aspergillus* spp. In this study, we have investigated specifically expressed TFs in the conidia of three *Aspergillus* species and constructed a gene knockout collection in the model organism *A. nidulans*. This collection provided an opportunity to explore the roles of 24 TFs in conidiogenesis and conidial properties, and we found a novel TF that is essential for proper conidial physiology. We explored, in detail, the functions of a spore-specific C_2_H_2_ zinc finger TF, *AN5003* (*sscA*), in conidial properties. The deficiency of *sscA* resulted in pleiotropic phenotypes, including impaired asexual spore formation and conidial long-term viability and abnormal germination. Furthermore, transcriptomic analyses revealed that the absence of *sscA* contributed to decreased stress response, transformations in cell wall integrity, and abnormal metabolism activities in *A. nidulans* **(Figure 9)**.

**Figure 9.**
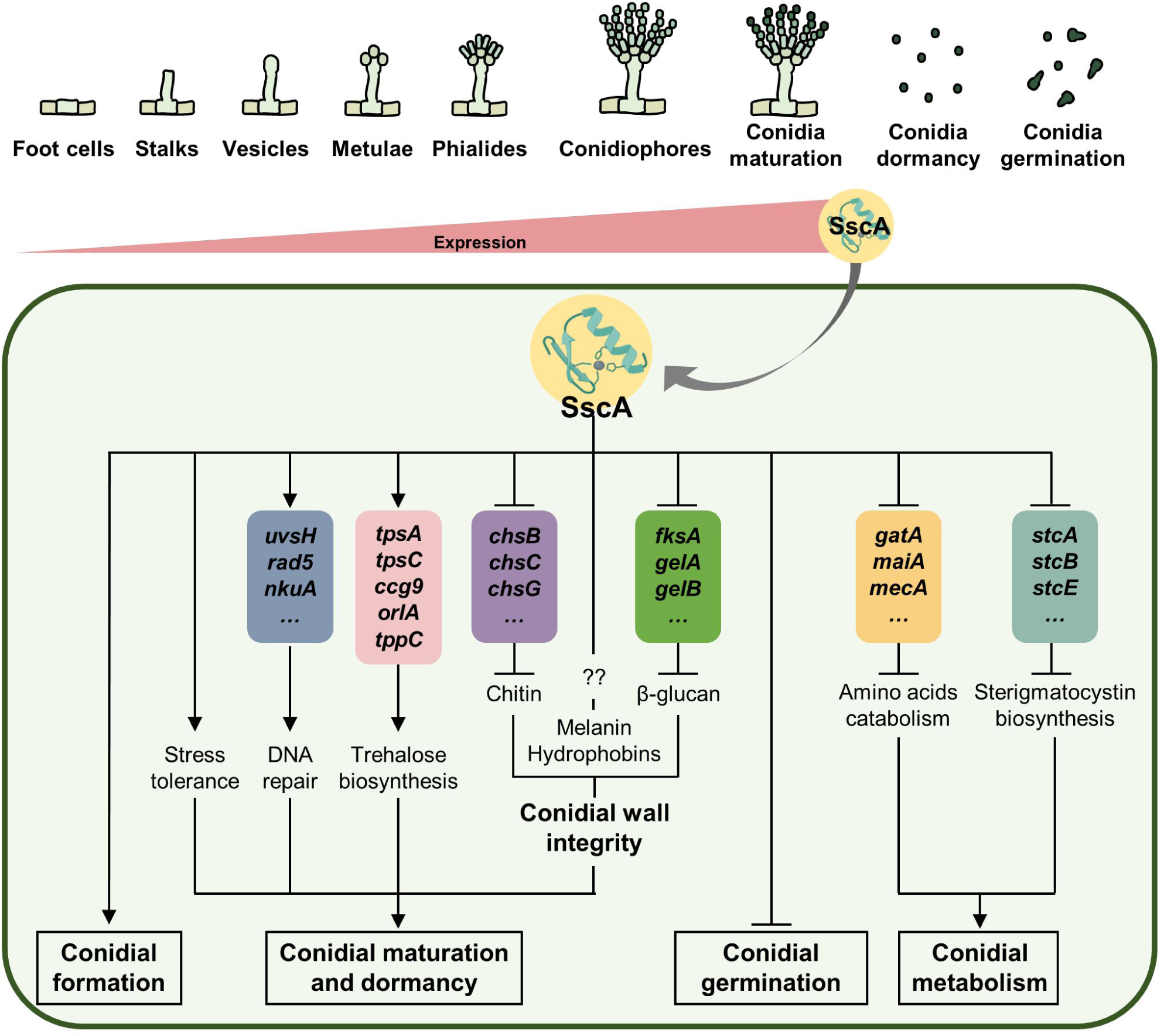
A proposed model depicting the functions of SscA in *A. nidulans* conidial formation, maturation, dormancy, and germination.

A preliminary clue into the functions of SscA in fungal physiology was obtained from the result that the absence of *sscA* led to formation of abnormal conidiophores with apricot-colored conidia. And the conidial production was significantly decreased in the absence of *sscA* **(Figure 3C-3D)**. Based on these results, we propose that an appropriate expression of SscA is required for proper asexual development in the lifespan of *A. nidulans*. SscA is conserved in eukaryotic microbes as well as *Aspergillus* species **(Figure 3A–3B)**, and the homologs of SscA also contribute to pleiotropic fungal physiology. In *C. albicans*, Bcr1 (biofilm and cell wall regulator 1) affects morphogenesis by differently expressing between yeast and hyphal form (48, 49). In *C. neoformans*, the absence of *usv101* influences the production of melanin (39, 50). In addition to its role in yeast, SscA plays important roles in the physiology of filamentous fungi. In *F. graminearum*, Nsf1 (nutrient and stress factor 1) is highly expressed in conidia compared to that in mycelium similar to SscA in *A. nidulans* (51). The disruption of *Fg*Nsf1 causes inhibited hyphal growth, defective asexual development, and fungal pigmentation in *F. graminearum.* Furthermore, Gcf6 (growth and conidiation regulatory factor 6) contributes to fungal growth, conidiation, and conidial germination in *Magnaporthe oryzae* (52). In addition, we have shown in the present study that *Afl*sscA and *Afu*sscA exerted conserved roles in asexual development and conidial physiology in *A. nidulans* **(Figure 8)**. Nevertheless, it is necessary to investigate whether SscA commonly contributes to fungal development in the plant pathogen *A. flavus* and the human pathogenic fungus *A. fumigatus* in future research.

As shown in **Figure S3**, mRNA levels of *sscA* increase during asexual development, implying that they might be regulated by central regulators of asexual development (BrlA or AbaA). We analyzed the *sscA* promoter region and found three BREs (BrlA-response element, 5′-(C/A)RAGGGR-3′) and one ARE (AbaA-response element, 5′-CATTCY-3′). Previous studies also suggested that Bcr1 is activated by Tec1 (homolog of AbaA) and regulates cell-surface-associated genes in *C. albicans* (49). Based on our results, we hypothesize that SscA expression is activated by AbaA, causing a gradual increase in mRNA levels of SscA following asexual development and asexual spore formation. To elucidate the genetic relationship between *abaA* and *sscA*, we measured levels of the *sscA* transcript in the asexual developmental stages of WT and the Δ*abaA* mutant. As shown in **Figure S8**, mRNA levels of *sscA* are considerably decreased in the absence of *abaA*. Next, we further evaluated whether SscA is coordinated by WetA, VosA, and VelB in *A. nidulans*. Our results have revealed that SscA is essential for normal conidial viability **(Figure 4A–4C)** and WetA, VosA, and VelB are the primary regulators of conidial maturation in *A. nidulans* conidia (42). Hence, we analyzed the multiomics data published in 2021 and confirmed that the transcript levels of *sscA* are not affected in the Δ*wetA,* Δ*vosA,* and Δ*velB* conidia. Overall, these results suggest that SscA expression is dependent on AbaA, but not on WetA, VosA, and VelB, for the proper asexual development of *A. nidulans*.

Our RNA-seq data showed that SscA is closely involved in the response to stresses and chemicals **(Figure 4A).** Also, we observed that the deletion of *sscA* led to lower trehalose contents and reduced resistance to UV, thermal, and oxidative stresses in conidia **(Figure 5D)**. Namely, SscA is essential for stress tolerance in the conidia of *A. nidulans*. Previous study has demonstrated that *usv1* null mutant is more sensitive to osmotic stress in *S. cerevisiae* (*53*). In *C. albicans*, the sensitivity of Δ*bcr1* to caffeine, lithium chloride, and copper is higher than that of WT (40). In *C. neoformans*, the deletion of *usv101* causes sensitivity to hydrogen peroxide, tert-butyl hydroperoxide, menadione, DTT, and AMB (39). However, the absence of *Fgnsf1* causes resistance to osmotic, oxidative, and metal cation stresses in *F. graminearum* (54). Collectively, the functions of SscA homolog in the stress resistance are species-specific.

Phenotypic and transcriptomic analyses of the *sscA* mutant have revealed that SscA is important for controlling ST biosynthesis **(Figure 6)**. Moreover, our analysis of the transcripts of the gene clusters of 23 secondary metabolites has revealed that SscA negatively regulates the production of other secondary metabolites in *A. nidulans* conidia. A previous study showed that *F*gNsf1 is not required for the production of the toxin zearalenone (40) but is required for the biosynthesis of the trichothecene mycotoxin, deoxynivalenol in *F. graminearum* (*54*). Therefore, the functions of SscA in secondary metabolism may be species-specific.

Genome-wide expression analyses have revealed that SscA affects cellular amino acid catabolic process in *A. nidulans* conidia. In fact, most genes related to amino acid catabolism in the Δ*sscA* mutant are highly expressed compared to that in WT strain, and consequently, the abundance of amino acids in the conidia of the Δ*sscA* mutant is less than those of WT **(Figure 7)**. Although it is not yet clear how these metabolic changes affect fungal development and conidial physiology, previous studies have reported that the amount of cellular amino acids was related to cell longevity in *Saccharomyces cerevisiae* (55, 56). Based on our result that the deletion of *sscA* causes deficient long-term spore viability, we suggest that SscA regulates spore longevity in part by coordinating the metabolism of amino acids.

The Cys_2_His_2_ Zinc finger protein is one of the largest families of TFs in eukaryotes and the secondly largest TFs in *A. nidulans* genome (57, 58). This type TF is commonly composed of two or three β-sheets and one α-helix, enabling specific DNA –binding. As trans-acting factors, C_2_H_2_ TFs regulate certain gene expression related to cellular development, differentiation, stress response, and metabolism in eukaryotes. Previous studies have demonstrated that C_2_H_2_ proteins also have pleiotropic effects on *A. nidulans* development, stress response, and metabolism: AslA, BrlA, FlbC, NsdC, SteA, NrdA, CreA and MtfA. BrlA is a central regulator of asexual development; AslA and FlbC also affect asexual development by activating *brlA* expression (59, 60). NsdC and SteA regulate normal sexual development (61). NrdA is required for oxidative stress resistance as well as normal sexual development (62). MtfA plays important roles for proper asexual and sexual development and secondary metabolism in *A. nidulans* (63). CreA is a well-known C_2_H_2_ TF for carbon catabolite repression, which regulates expression of various hydrolytic enzymes (64). In this study, we characterize a novel C_2_H_2_-type Zinc finger TF in *A. nidulans*. SscA has one Cys_2_His_2_ zinc finger domain composed of two β-sheets and one α-helix, patterned as C-X_5_-C-X_12_-H-X_3_-H. Like other C_2_H_2_ TF, SscA also act as a multifunctional regulator for asexual spore formation, conidial stress tolerance, and metabolism. Zinc-binding proteins can act as activators, repressors, or both activators and repressors for certain genes. When considering the number of upregulated (2,695) and downregulated (1,874) genes in deletion of *sscA*, SscA might function primarily as a transcriptional repressor. However, further studies are required to understand the molecular functions of SscA as a TF.

To summarize, we investigated the DEGs between conidia and hyphae and generated a collection of TFs that are highly expressed in the conidia of three *Aspergillus* species. Through various phenotypic analyses, we identified a novel Cys_2_His_2_ TF, SscA, in *A. nidulans*. Based on our in-depth studies, we conclude that SscA plays an essential role in the regulation of conidial formation, maturation, dormancy, germination, and metabolism in *A. nidulans*. Our results also suggest that the functions of *Afl*sscA and *Afu*sscA are conserved in *A. nidulans* conidia.

## Materials and methods

### Strains, media, and culture conditions

*Aspergillus* strains used in this study are listed in **Table S5**. *A. nidulans* strains were cultured in liquid or solid minimal medium supplemented with 1% glucose (MM) (65). *A. flavus* and *A. fumigatus* strains were grown on MM with 0.1% yeast extract (MMYE) (31). For plasmid manipulation, *Escherichia coli* DH5α was grown in Luria–Bertani medium (BD, Difco, Franklin Lakes, NJ, USA) containing ampicillin (100 μg/mL) (Sigma-Aldrich, St. Louis, MO, USA).

### Generation of TF deletion mutants

Double-joint PCR (DJ-PCR) was used for the generation of the TF deletion mutant collection, and oligonucleotides used in this study are described in **Table S6** (43). Briefly, the 5ʹ- and 3ʹ-flank regions of each TF were amplified using the primer pairs 5ʹ DF/3ʹ tail and 5ʹ tail/3ʹ DR of each TF gene, respectively, with *A. nidulans* FGSC4 genomic DNA (gDNA) as a template. The *A. fumigatus pyrG* marker was amplified from *A. fumigatus* Af293 gDNA using the primer set OHS1542/OHS1543. Three fragments, including 5ʹ-flank, 3ʹ-flank, and *pyrG*^+^, were fused through second and third PCR with the primer set 5ʹ NF/3ʹNR of each TF gene, and the deletion cassette was transformed into the *A. nidulans* strain RJMP 1.59.

### Construction of *sscA*-complemented strains

For constructing the *sscA*-complemented strain in *A. nidulans*, the *AnisscA* gene region with its promoter was amplified using the primer pair OHS1711/OHS1713, digested with *Not*I, and cloned into pHS13 (26). The resulting plasmid pYE7.1 was then introduced into the recipient Δ*AnisscA* strain TSH31.1 to produce TYE51.1∼2.

For the complementation of the Δ*AnisscA* strain with *A. flavus*, the ORF, and its upstream promoter region were amplified using *A. flavus* NRRL 3357 gDNA and the primer set OHS1876/ OHS1780, digested with *Not*I, and cloned into pHS13. The resulting plasmid pYE9.1 was transformed into the recipient Δ*AnisscA* strain TSH31.1 to produce TYE57.1∼2.

For the complementation of the Δ*AnisscA* strain with *A. fumigatus*, the ORF, and its upstream promoter region were amplified using *A. fumigatus* Af293 gDNA and the primer set OHS1763/OHS1764, digested with *Not*I, and cloned into pHS13. The resulting plasmid pYE10.1 was transformed into the recipient Δ*AnisscA* strain TSH31.1 to produce TYE58.1. All the complemented strains were selected among the transformants and screened by PCR and quantitative reverse transcription-PCR (qRT-PCR).

### Transcriptomic analyses and GO analyses

For transcriptomic analyses of hyphae, the conidia (10^6^/mL) of three *Aspergillus* species were inoculated into 100 mL of liquid MM or MMYE and incubated at 37°C for 14 h. The cultured samples were filtered, washed, squeeze-dried, and stored at −80°C. For transcriptomic analyses of conidia, 2-day-grown conidia were collected and filtered through Miracloth (Calbiochem, San Diego, CA, USA) (66).

Total RNA isolation was performed as previously described (67). Each sample was homogenized using a Mini-Bead beater (BioSpec Products Inc., Bartlesville, OK, USA) with 0.3 mL of glass beads (Daihan Scientific, Wonju, South Korea) and 1 mL of TRIzol reagent (Invitrogen Waltham, MA, USA) and centrifuged. The supernatant was mixed with cold isopropanol and centrifuged. The RNA pellets were washed with 70% ethanol and then dissolved in DEPC-DW (Bioneer, Daejeon, South Korea). The extracted RNA samples were treated with DNase I (Promega, Madison, WI, USA) and purified using the RNeasy Mini Kit (Qiagen, Hilden, Germany), after which library construction and Illumina platform sequencing were performed by Theragen Bio (Suwon, South Korea). Briefly, for paired-end sequencing, RNA integrity was evaluated on an Agilent 2100 Bioanalyzer system (Agilent Technologies, Santa Clara, CA, USA), and the RNA library was constructed using the TruSeq Stranded mRNA Sample Prep Kit (Illumina, San Diego, CA, USA). Each sample with sequencing adaptors was sequenced in a single lane on the Illumina Novaseq 6000 system. Recorded Fast files were analyzed with FastQC and trimmed using Trimmomatic. Filtered reads were mapped to the *A. nidulans* FGSC A4, *A. flavus* NRRL3357, and *A. fumgiatus* Af293 transcriptome using the aligner STAR v.2.3.0e software. Based on read counts, the DEGs were identified, and normalized using the DESeq2 method. All RNA-seq experiments were performed with at least two biological replicates.

For the characterization of DEGs, GO analyses were conducted using R package and FungiDB (https://fungidb.org/fungidb/app). DEGs, which were identified based on two-fold change and *p* value < 0.05, were subjected to Fisher’s exact test, and all processed data were plotted using R studio (v. 3.6.3).

### Quantitative real-time PCR

For the induction of asexual development, cultured mycelia were transferred to solid MM and incubated at 37°C for indicated time points under light condition (68). From the isolated total RNA, cDNA was synthesized using GoScript Reverse transcriptase (Promega, Madison, WI, USA), and real-time PCR was performed using iTaq Universal SYBR Green Supermix (Bio-Rad, Hercules, CA, USA) on CFX96 Touch Real-Time PCR (Bio-Rad) (67). The primers used for qRT-PCR are listed in **Table S6**. The fold expression was computed using the 2^-ΔΔCt^ method, and the expression was normalized to that of β-actin, which was used as a control.

### Conidial production, viability, and germination assay

To quantify the production of conidia, each strain was point-inoculated onto solid MM media and incubated at 37°C for 5 days. Then, the conidia were harvested using distilled water containing 0.02% Triton X-100 and counted using a hematocytometer under a Leica DM500 microscope (Leica, Wetzlar, Germany).

For spore viability assay, conidia were grown for 2 or 10 days and collected using DW containing 0.02% Triton X-100. Then, approximately 1 × 10^2^ conidia were spread onto solid MM and incubated for 2 days at 37°C. The viability of conidia was calculated as the ratio of the number of viable colonies from 10-day-old conidia to the number of viable colonies from 2-day-old conidia in triplicate (31, 66).

For the assay of conidial germination, approximately 10^7^ conidia grown for 2 days were spread onto MM agar with or without glucose and cultured at 37°C. The germination rate was calculated as the ratio of the number of germinated spores to that of total spores every hour (67).

### Spore trehalose, stress tolerance, and β-glucan assay

For the assay of spore trehalose content, conidia (2 × 10^8^) from 2-day-old cultures were collected, resuspended in ddH_2_O, and incubated at 95°C for 20 min. The supernatant was mixed with an equal volume of 0.2 M sodium citrate (pH 5.5) and incubated at 37°C for 8 h with or without (as a negative control) trehalase (Sigma). The amount of glucose generated from trehalose was assayed using a Glucose Assay Kit (Sigma) in triplicate (31, 68).

For evaluating the spore tolerance against thermal and oxidative stresses, approximately 10^3^ conidia were incubated at 37°C in 0.1 M H_2_O_2_ for 30 min. Then, the samples were diluted, spread onto solid MM, and incubated for 2 days at 37°C. For the conidial UV stress test, approximately 10^2^ conidia were spread onto solid MM and UV-irradiated using a UV crosslinker (Spectrolinke XL-1000 UV crosslinker, Thomas Scientific, Swedesboro, NJ, USA). Then, the plates were incubated for 2 days at 37°C. The survival rates were calculated as the ratio of the number of viable colonies to that of the untreated control (67, 69).

To examine β-1,3-glucan analysis, approximately 10^3^–10^4^ conidia grown for 2 days were resuspended in 25 μL, mixed with Glucatell^®^ reagent, and incubated at 37°C for 30 min. Then, the reaction was stopped using diazo coupling reagents, and the optical density of the samples was read at 540 nm (41).

### Transmission electron microscopy

TEM analysis was performed with a modification of a previously described method (41). Briefly, 10-day-old conidia (5 × 10^8^) were collected and fixed in Karnovsky’s fixative at 4°C overnight. The samples were then fixed with 1% osmium tetraoxide (Sigma), dehydrated in graded ethanol, and resuspended in propylene oxide (Sigma) and poly/bed^®^ 812 (Polyscience, Warrington, PA, USA), sequentially. For embedding, the samples were added to poly/bed^®^ 812 with 2% DMP-30 and solidified in an oven (65°C) overnight. Polymerized samples were sectioned on a Leica EM UC7 cryo-ultramicrotome equipped with a diamond knife and stained with uranyless (EMS, Hatfield, PA, USA) and lead citrate (EMS). The stained sections were viewed under a Hitachi HT7700 transmission electron microscope (Hitachi, Soto-Kanda, Chiyoda-ku, Tokyo).

### Amino acid analysis in conidia

For quantifying the amount of amino acids in conidia, approximately 2 × 10^9^ conidia grown for 2 days were prepared, mixed with 0.5 mL of HPLC-grade acetonitrile-methanol-water (40:40:20, v/v/v) and 0.3 mL glass beads, and homogenized using a Mini-Bead beater (BioSpec Products Inc.) (42). The lysates were centrifuged, and the supernatant was filtered using a 0.45-μm hydrophobic PTFE filter (Falcon, Corning, NY, USA). The purified samples were pretreated with methanol:acetonitrile:DW (2:2:1, v/v/v) and analyzed using the amino acid analyzer L-8900 (Hitachi) consisting of 4.6 mm ID × 60 mm column packed with ion exchange resin. The amount of amino acids was measured using the colorimetric ninhydrin method (Wako, Tokyo, Japan) and calculated using the mixture of amino acid standard solution type ANII and type B (Wako).

### Sterigmatocystin production analysis in conidia

Sterigmatocystin (ST) was extracted from conidia as described previously (67), with some modifications. Approximately 2 × 10^8^ conidia were homogenized using a Mini-Bead beater (BioSpec Products Inc.) with 0.3 mL of glass beads (Daihan Scientific) and 1 mL of chloroform (Sigma). After centrifugation, the organic phase was mixed with DW for refining secondary metabolites and centrifuged. The purified organic phase was transferred to a new glass vial and dried in an oven (65°C) overnight. The samples were resuspended with 0.1 mL of chloroform, spotted in Thin-layer chromatography (TLC) silica plate coated with the flourescent indicator F254 (Merck Millipore, Burlington, MA, USA), and resolved in TLC plates containing toluene:ethyl acetate:acetic acid (8:1:1, v/v/v). The Thin-layer chromatography (TLC) plates were treated with 1% aluminum hydroxide hydrate (Sigma), and images were captured under UV exposure (366 nm). The relative intensities of ST were calculated using the ImageJ software.

### Microscopy

Photographs of colonies were taken using a Pentax MX-1 digital camera. Photomicrographs were taken using a Leica DM500 microscope equipped with Leica ICC50 E and Leica Application Suite X software.

### Statistical analysis

Statistical differences between the control and target strains were evaluated using Student’s unpaired *t*-test. Data are reported as mean ± standard deviation. *P* values <0.05 were considered significant.

### Data availability

All RNA-seq data files are available at the NCBI Sequence Read Archive (SRA) database under the accession number PRJNA967141, PRJNA967140, and PRJNA967143; for conidia and hyphae of three *Aspergillus* RNA-seq and PRJNA836364; for WT and *sscA* of *A. nidulans* conidia RNA-seq.).

## Acknowledgments

The work by HSP was supported by the National Research Foundation of Korea (NRF) grant to HSP funded by the Korean government (MSIP: 2020R1C1C1004473).

The work by SYE was supported by the National Research Foundation of Korea (NRF) grant to HSP funded by the Korean government (MSIP: 2021R1A6A3A13044577).

This research was supported by a project to train professional personnel in biological materials by the Ministry of Environment and the Korea Basic Science Institute (National Research Facilities and Equipment Center) grant funded by the Ministry of Education (2021R1A6C101A416).

